# The critical role of BTRC in hepatic steatosis as an ATGL E3 ligase

**DOI:** 10.1101/2022.11.15.516629

**Authors:** Weiwei Qi, Zhenzhen Fang, Chuanghua Luo, Honghai Hong, Yanlan Long, Zhiyu Dai, Junxi Liu, Yongcheng Zeng, Ti Zhou, Yong Xia, Xia Yang, Guoquan Gao

## Abstract

**Objective:** Non-alcoholic fatty liver disease (NAFLD), characterized by hepatic steatosis, is one of the most common causes of liver dysfunction. ATGL is closely related to hepatic steatosis as the speed-limited triacylglycerol lipase. Nevertheless, the expression and regulation of ATGL in NAFLD remain unclear.

**Methods:** Using immunohistochemistry and qRT-PCR to detect the expression of ATGL and BTRC in different models with hepatic steatosi**s**. Co-IP evaluated the binding of ATGL and BTRC. Knockdown of BTRC employed by adenoviruses and then analyzed the ATGL expression, triglyceride levels, and lipid droplets accumulation.

**Results:** Our results revealed that ATGL protein level was decreased in animal and cellular models of hepatic steatosis and the liver tissues of cholangioma/hepatic carcinoma patients with hepatic steatosis, while the ATGL mRNA level had hardly changed; which means the decreased ATGL mainly degraded through the proteasome pathway. BTRC was identified as the E3 ligase for ATGL, up-regulated, and negatively correlated with ATGL level. Moreover, adenovirus-mediated knockdown of BTRC ameliorated hepatic steatosis *via* up-regulating ATGL level.

**Conclusions:** Our study demonstrates a crucial role of elevated BTRC in hepatic steatosis through promoting ATGL proteasomal degradation as a new ATGL E3 ligase and suggests BTRC may serve as a potential therapeutic target for NAFLD.

**Funding:** This study was supported by The National Natural Science Foundation of China (Grants 82070888, 82070882, 82100917, 81872165, 82273116, 82203661, 81901557 and 81902693); Key Project of Nature Science Foundation of Guangdong Province, China (Grant 2019B1515120077); National Key R&D Program of China (Grant 2018YFA0800403); Guangdong Special Support Program for Young Top Scientist (Grant 201629046); Guangdong Natural Science Fund (Grant 2019A1515011810, 2021A1515010434, 2022A1515012423 and 2022A1515012513); Key Sci-Tech Research Project of Guangzhou Municipality (Grants 202201010820) Fundamental Research Funds for the Central Universities (Grant 50000-31620106); China Postdoctoral Science Foundation (Grant 2021M703679, 2020M683110).

## Introduction

Non-alcoholic fatty liver disease (NAFLD), characterized by hepatic steatosis and lipid storage, is currently the most frequent global liver disease^1^. Disease presentation includes various clinical phenotypes ranging from hepatic steatosis (non-alcoholic fatty liver) to non-alcoholic steatohepatitis (NASH), which can progress to liver fibrosis, cirrhosis, and hepatocellular carcinoma^2^. NAFLD increases the risk of type 2 diabetes mellitus (T2DM), cardiovascular (CVD) and cardiac diseases, and chronic kidney disease (CKD)^3^. Additionally, recent studies have shown that NAFLD was associated with a 93% higher relative risk of overall mortality, and a 20-year absolute excess risk of 15.3%, primarily due to increased cancer- and cirrhosis-specific mortality^4^. Therefore, simple steatosis can no longer be ignored by medicine, and it is essential to explore its occurrence’s detailed molecular mechanism.

The imbalance between the source and metabolic pathway of lipid, including defective lipolysis, decreased lipid export, excessive free fatty acids (FFAs) uptake, and abnormally enhanced *de novo* fatty acids synthesis, may contribute to the development of hepatic steatosis^5,6^. Hydrolysis of triacylglycerol (TAG) to generate FFAs and glycerol require three consecutive steps that involve at least three kinds of enzymes: Adipose triglyceride lipase (ATGL) initiates the lipolysis process, converting TAGs to diacylglycerols (DGs); hormone-sensitive triglyceride lipase (HSL) is mainly responsible for the hydrolysis of DGs to monoacylglycerols (MGs), which can be further hydrolyzed by MG lipase (MGL)^7^. ATGL is the latest member of the lipolytic enzyme, first described in 2004, and belongs to the family of patatin domain-containing proteins, including nine human and eight murine members^8,9^. ATGL knockout animals had enlarged fat depots and drastically reduced TAG hydrolase activities in white adipose tissue (WAT) lysates^10^. Condition deletion of liver ATGL in mice revealed progressive hepatic steatosis and enhanced lipid droplet (LD) accumulation^11^. Moreover, NAFLD patients with insulin resistance had higher liver steatosis grades and lower ATGL expression^12^. These studies underscored the importance of lipolysis and ATGL levels to lipid homeostasis in the liver. To date, most studies on hepatic ATGL were focused on its enzymes’ activity and transcriptional level regulation^13,14^. However, little is known about the post-transcriptional regulation of ATGL and how this process contributes to the development of hepatic steatosis.

Ubiquitin-dependent proteasome degradation pathway plays a pivotal role in cellular protein turnover^15^. Ubiquitin-dependent proteolysis of proteins impacts various cellular processes such as cell cycle progression, transcription, antigen presentation, receptor endocytosis, fate determination, and signal transduction^16^. Stepwise ubiquitination of a target protein is achieved through three enzymes: E1 ubiquitin (Ub) activating enzymes, E2 ubiquitin conjugating enzymes, and E3 ubiquitin ligases^17,18^. BTRC, also called β transducin repeat containing proteins (β-TrCP), is a member of the F-box family of proteins and serves as a substrate recognition component of E3 ubiquitin ligases^19^. It has been showed that BTRC could recognize a wide range of cellular targets for degradation to regulate multiple biological processes, including cell cycle regulators Wee1 and Cdc25A, negative regulator of NF-κB signaling IκBα and other important signal transduction molecules like β-catenin, Snail^20^. However, it is not known whether BTRC could act as the E3 ligase of ATGL for degradation and be involved in lipid metabolism and hepatic steatosis.

In this study, we demonstrated the role of BTRC in ATGL degradation and hepatic steatosis. Moreover, we proved the hypothesis that knockdown of BTRC could increase ATGL levels, inhibit TAG accumulation and ameliorate hepatic steatosis. This study suggests that BTRC might be a novel target for the treatment of NAFLD.

## Materials and Methods

### Antibodies and reagents

Rabbit anti-ATGL antibody, mouse anti-Ub antibody, and rabbit anti-BTRC antibodies were from Cell Signaling Technology (Boston, MA); the DAPI and β-actin antibody were from thermo scientific; MG-132 was from Merck (Catalog number 47490, Merck, Darmstadt, Germany); Lysosomal inhibitor ammonium chloride (NH_4_Cl) was from Sigma-Aldrich (Catalog number 254134, Sigma-Aldrich, St. Louis, MO).

### Cell Culture and Transfection

Human hepatoma cell line HepG2, human embryonic kidney cell line HEK293A (obtained from ATCC), and human liver cell line Chang liver(obtained from the cell bank of the Chinese Academy of Sciences in Shanghai)were cultured in high glucose DMEM supplemented with 10% FBS containing 1% penicillin/streptomycin. Primary hepatocytes were purified from the liver tissue of mice by perfusing the liver with 0.4 mg/mL collagenase IV (Catalog number C5138, Sigma, USA) to digest hepatocytes and then cultured in Willian’s E medium with hepatocyte maintenance culture supplement containing 10% FBS and 1% penicillin/streptomycin. Transient transfections with plasmids and small interfering siRNA were done using Lipofectamine 2000 reagent (Catalog number 11668027, Thermo Fisher Scientific, Waltham, MA) according to the manufacturer’s instruction.

### Plasmids and Vectors

Human BTRC, FBW7, FBW5, and ATGL clones were constructed using Ruyilian Kit from SiDanSai Biotechnology (Shanghai, China) according to the manufacturer’s instruction. siRNA for BTRC was purchased from Ribobio Company (Guangzhou, China). Mutagenesis of the ATGL clone was done using In-Fusion technology from Clontech Company (Catalog 639650, Clontech, Japan). Details of primers and cloning construction steps for plasmid and mutagenesis were in **Table S1** and **Table S2**.

### Animal Experiments

Male C57BL/6J mice (7-8 weeks old) were from Vital River (Beijing, China). The mice were housed under specific pathogen free (SPF) conditions with a 12-hours light/dark cycle, a controlled humidity (40%-70%), and a stable temperature (22±3 °C). All animal experiments have been reviewed and approved by the Animal Ethics Committee of Sun Yat-sen University. The use and handling of animals were performed strictly in accordance with the Guidelines for the Use of Laboratory Animals by Sun Yat-sen University. Animals were fed a chow or high fat diet (HFD, Research Diets, D12492). The adenoviruses for the knockdown of BTRC expression and the control adenoviruses were from Obio (Shanghai, China). To verify the interference effect, the viruses were used to infect hepa1-6 cells at an MOI of 10. Viruses were diluted in PBS and administered through tail vein injection for animal experiments. Mice were injected with 1×10 ^9^ plaque forming units of adenoviruses.

Male naturally obese C57BL/6J mice (6 months old) were from Vital River (Beijing, China), which were all fed on a regular diet, and the most obese mice were selected for the hepatic steatosis group(38.16±0.12g), and the mice with average weight were selected for the control group(30.44±0.44g).

### Measurement of triglyceride, FFA, and glycerol levels

50 mg of liver tissue was homogenized, and lipids were extracted from the liver and cultured hepatocytes in chloroform/methanol (2:1 v/v) as described ^21^. The amount of triglycerides, glycerol and FFA were measured by using EnzyChrom™ triglyceride assay kit (Bioassay Systems), glycerol assay kit (Bioassay Systems), and free fatty acid assay kit (Bioassay Systems), respectively.

### Histological analysis of human tissues

The surgical liver specimens of cholangioma patients (from The First Affiliated Hospital of Sun Yat-sen University) and hepatocellular patients (from The Second Affiliated Hospital of Sun Yat-sen University) with hepatic steatosis or non-hepatic steatosis, and the liver samples of naturally obese and high fat diet mouse were fixed with 4% (w/v) paraformaldehyde overnight. Sections (5-*μ*m) were prepared from the paraffin-embedded tissues and were stained for ATGL and BTRC, respectively. Detection was performed using ABC HRP Kit (Vector Laboratories) with DAB followed by counterstaining with hematoxylin or methyl green. Images were obtained using a light microscope (Nikon Corporation, Tokyo, Japan). All samples were collected with patients’ consent, and the experiments were approved by the ethics committee of Sun Yat-sen University (Guangzhou, China).

### qRT-PCR

Total RNA was isolated using Trizol (Invitrogen) and reverse-transcribed (Takara). BTRC was amplified with the primers of 5’-GGAGAAGACTTTGACCAGCG-3’ and 5’-CTTTGGAATTCGAGTCGAGC-3’; FB5 was amplified with the primers of 5’-GGAGAAGACTTTGACCAGCG-3’ and 5’-CTTTGGAATTCGAGTCGAGC-3’; FB7 was amplified with the primers of 5’-GGAGAAGACTTTGACCAGCG-3’ and 5’-CTTTGGAATTCGAGTCGAGC-3’; Amplification reactions were performed with an initial denaturation step at 94°C for 2 min, followed by 40 cycles consi sting of adenaturation step at 94°C for 5 min, an annealing step at 55°C for 30 s, and an extension step at 72°C for 2 min.

### Co-immunoprecipitation

For immunoprecipitations, 1000μg cell lysates were incubated with 8μg of anti-ATGL or anti-BTRC antibody overnight at 4°C, followed by the addition 40μl of Protein A or G–Sepharose beads (Calbiochem) and incubated for 4h at 4°C. Immunoprecipitates were washed four times with lysis buffer, eluted with loading buffer, and performed Western Blot Analysis.

### Oleic acid-induced hepatic steatosis cell model

HepG2 cells were treated with indicated concentration OA solution (Sigma-Aldrich) and cultured in Dulbecco’s modified Eagle’s medium (Hyclone; GE Healthcare Life Sciences, USA) with 0.2% fetal bovine serum. After a different time, the medium was removed, and the cells were harvested for Western blot or Oil Red O Staining.

### Oil Red O Staining

Oil Red O staining was performed as described before ^22^. Photographs were taken with a camera under optimal illumination.

### Statistical Analysis

The Student unpaired two-tailed *t* test was used to evaluate the statistical significance between two groups, and ANOVA was used for more than two groups, followed by Fisher’s least significant difference test. Statistical significance was predefined as *P*<0.05.

## Results

### ATGL protein is down-regulated in hepatic steatosis

To assess the ATGL expression in hepatic steatosis, we collected surgical specimens of cholangioma patients with hepatic steatosis and non-hepatic steatosis and then performed immunohistochemistry staining. The result showed that ATGL was significantly down-regulated in cholangioma patients with hepatic steatosis **(Fig 1A)**. Similar results were found in paracancerous liver tissues of hepatic carcinoma patients with hepatic steatosis **(Fig 1B)**. Furthermore, we collected 6-months-old naturally obese mice with hepatic steatosis to exclude the effect of HFD, and found liver ATGL expression in the obesity group was significantly reduced compared with the control group **(Fig 1C)**. Moreover, the liver ATGL protein levels tended to decline in the hepatic steatosis mouse model induced by HFD for 12 weeks, and remarkably decreased in 16 weeks **(Fig 1D)**, but not for ATGL mRNA level **(Fig 1E)**. Next, we investigated hepatic steatosis patients’ data sets (GSE89632 and GSE48452), and found that the liver ATGL mRNA levels were not decreased in the patients with hepatic steatosis **(Fig 1F)**. In addition, we employed the hepatic steatosis cell model, which was treated with 50μM oleic acid *in vitro*. **Fig 1G** verified that the ATGL protein level was noticeably down-regulated after 15 days of OA treatment, while the ATGL mRNA level had no change **(Fig 1H)**. Taken together, these results indicate that the ATGL level is perceptibly reduced in hepatic steatosis, and post-transcriptional regulation may play a vital role in the decreased ATGL level.

**Figure 1.**
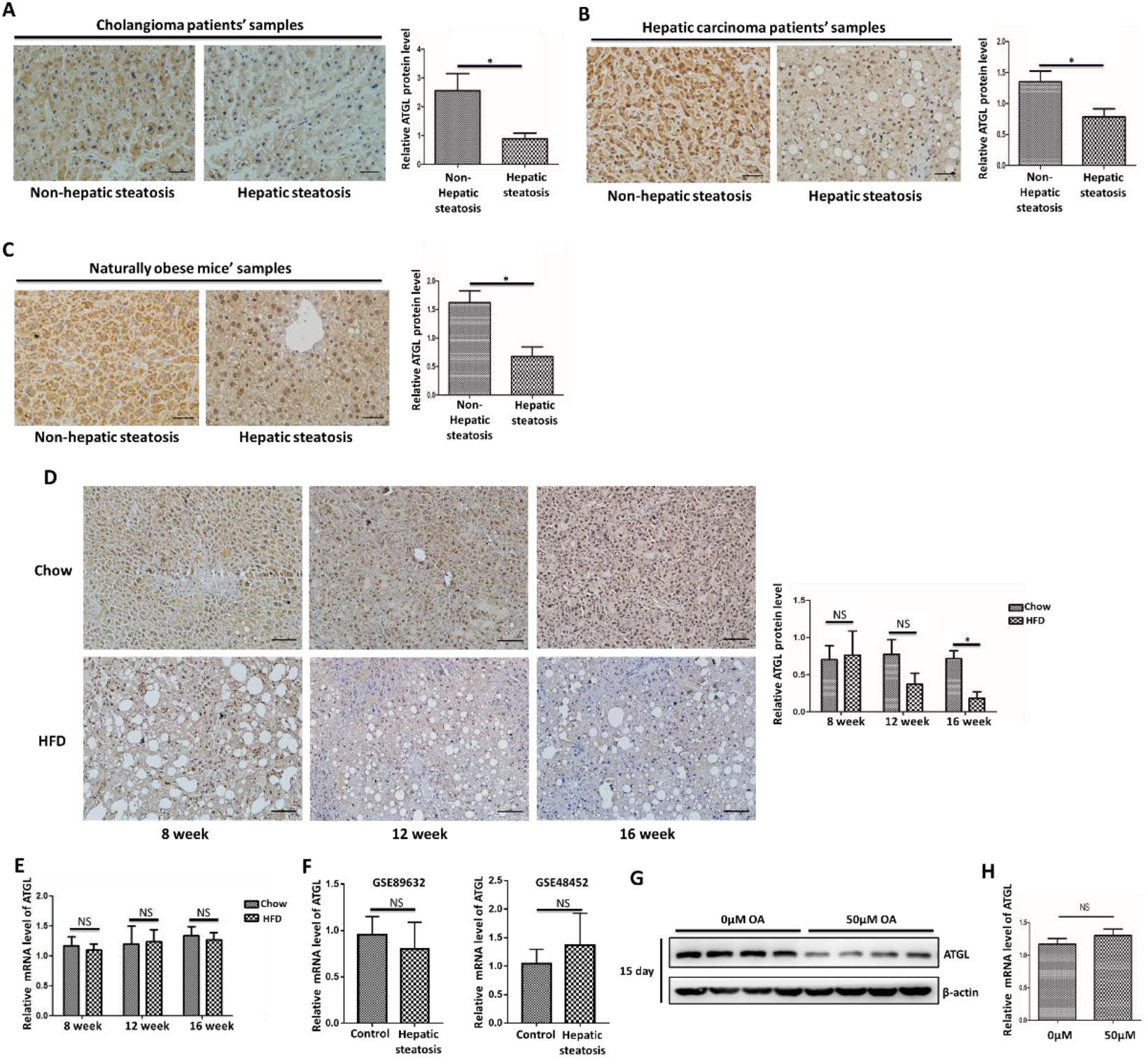
ATGL protein is down-regulated in hepatic steatosis. A-D ATGL protein expression level was analyzed by IHC images in liver of cholangioma patients with hepatic steatosis (A, n=3 samples/group), hepatic carcinoma patients with hepatic steatosis (B, n=11 samples/group), naturally obese mouse with hepatic steatosis (C, n=6 mice/group) and hepatic steatosis mouse model induced by HFD for 8 weeks, 12 weeks and 16 weeks (D, n=6 mice/group). E ATGL mRNA level was analyzed by qRT-PCR in hepatic steatosis mouse model induced by HFD for indicated times. F ATGL mRNA levels in the livers from patients with hepatic steatosis (Control, n=23; steatosis, n=18 for GEO accession number GSE89632. Control, n=12; steatosis, n=9 for GEO accession number GSE 48452) G-H HepG2 cells were treated with 50μM OA for 15 days. ATGL expression (G) was examined by immunoblotting and ATGL mRNA level (H) was detected by qRT-PCR. β-actin was used as a loading control. *Scale bars*: 50μm (A-C) and 100μm (D).

### ATGL is degraded through the proteasome pathway in liver

Post-transcriptional level regulation plays a crucial role in cellular protein turnover mainly through the proteasome and lysosome pathways^15^. To evaluate which pathway contributes to the decline of ATGL turnover in hepatic steatosis, we first treated hepatocytes with different protease inhibitors, lysosomal inhibitor ammonium chloride (NH_4_Cl), and proteasome specific inhibitor MG132. As shown in **Fig S1A** and **S1B**, there was no change of ATGL in Chang liver and HepG2 cells with the treatment of NH_4_Cl. Nevertheless, the amount of ATGL was significantly accumulated after treatment with the proteasome specific inhibitor MG132 in Chang liver, HepG2, and HEK293A cells **(Fig 2A, 2C&2E)**. To further confirm whether ATGL was degraded through the proteasome pathway, we treated hepatocytes with cycloheximide (CHX), which is described as a fungicide that inhibits the synthesis of proteins in eukaryotes. As shown in **Fig 2B and 2D**, ATGL was gradually decreased as time elonged under CHX treatment, but the degradation was blocked in the presence of MG132 and CHX. These results indicate that the proteasome-mediated degradation pathway regulates ATGL degradation in hepatocytes. In addition, the ATGL protein level in HepG2 cells was down-regulated when treated with OA for 12 days, which could be stalled after treatment with MG132 **(Fig 2F)**, while the ATGL transcriptional level was not affected **(Fig 2G)**. Overall, these results suggest that ATGL degeneration exists *via* proteasome pathway both in hepatocytes and hepatic steatosis conditions.

**Figure 2.**
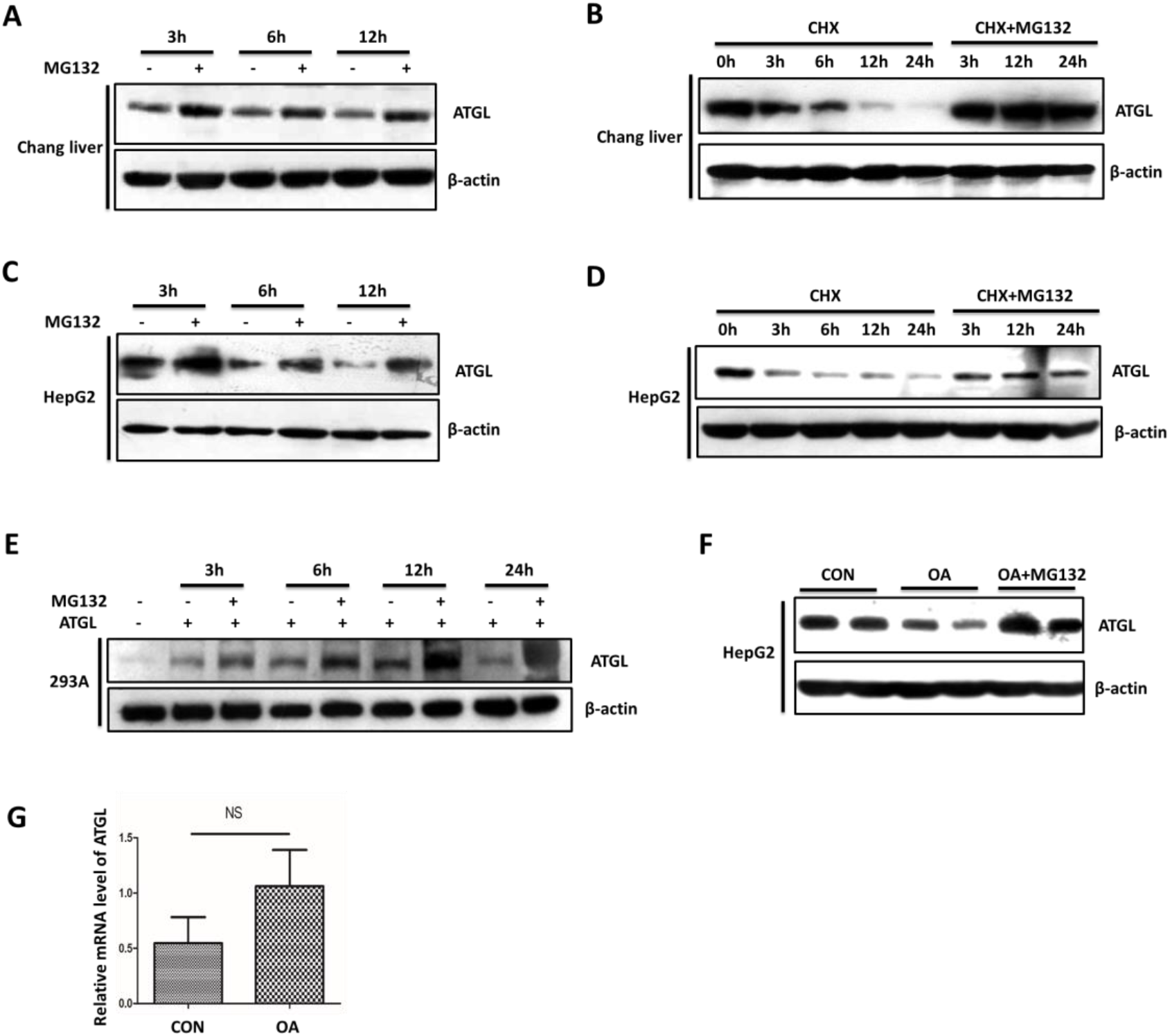
ATGL is degraded through proteasome pathway in hepatic steatosis. A-D ATGL was detected by immunoblotting in chang liver cells (A, B) and HepG2 cells (C, D), which were treated with cycloheximide(CHX) or CHX and MG132 for indicated time. E HEK293A cells were transiently transfected with ATGL plasmids for 24h, followed by treatment with/without MG132 for indicated time. ATGL expression was examined by immunoblotting. F, G Hepatic steatosis HepG2 cells model was induced by 50μM OA for 12 days followed by treating with MG132 for 24h and for western blot analysis and qRT-PCR (G). β-actin was used as a loading control. Data are represented of at least 3 independent experiments. CHX: 2μg/mL, MG132: 10μM.

### BTRC down-regulates the ATGL level and is negatively correlated with ATGL in hepatic steatosis

To identify which E3 ligase contributes to ATGL proteasome degradation, we first used bioinformatics to predict E3 ligase that maybe bind to ATGL. The STRING database (https://string-db.org/) indicated that RNF7 might be interacted with ATGL (predicted interaction scores=0.465, **Fig 3A**). RNF7 combined with F-box proteins to form the SCF (SKP1-CUL1-F-box protein) E3 ubiquitin ligase complex, which could mediate the ubiquitination and subsequent proteasomal degradation of target proteins ^23,24^. In addition, the STRING database also indicated that RNF7 combined with SKP1-CUL1-F-box protein to form the SCF complex **(Fig 3B)**. Therefore, we selected F-box protein FBW7, FBW5, and BTRC to evaluate their effects on ATGL protein level. As shown in **Fig 3C**, FBW5 and BTRC could down-regulated the ATGL protein level visibly, while FBW7 had no impact on ATGL, suggesting that FBW5 and BTRC may be ATGL’s E3 ligase. To determine which of FBW5 and BTRC is the real E3 ligase of ATGL in hepatic steatosis, we evaluated the expression of FBW5 and BTRC in the hepatic steatosis mouse model induced by HFD. The results showed that only the mRNA level of BTRC was elevated in hepatic steatosis, while the levels of FB5 did not change **(Fig 3D&E)**. Correspondingly, the BTRC protein was up-regulated in paracancerous liver tissues of hepatic carcinoma patients with hepatic steatosis and mouse models with hepatic steatosis induced by HFD, which was negatively correlated with the expression of ATGL protein **(Fig 4A&B)**. Interestingly, where the ATGL level was decreased, the level of BTRC was increased, and vice versa **(Fig 4C)**. Furthermore, in 15 days oleic acid-induced hepatic steatosis cell model, the level of BTRC was significantly elevated, while the ATGL level was markedly down-regulated **(Fig 4D)**. Taken together, these results suggest that BTRC may be the E3 ligase of ATGL in hepatic steatosis, which could regulate the proteasome pathway degradation of ATGL.

**Figure 3.**
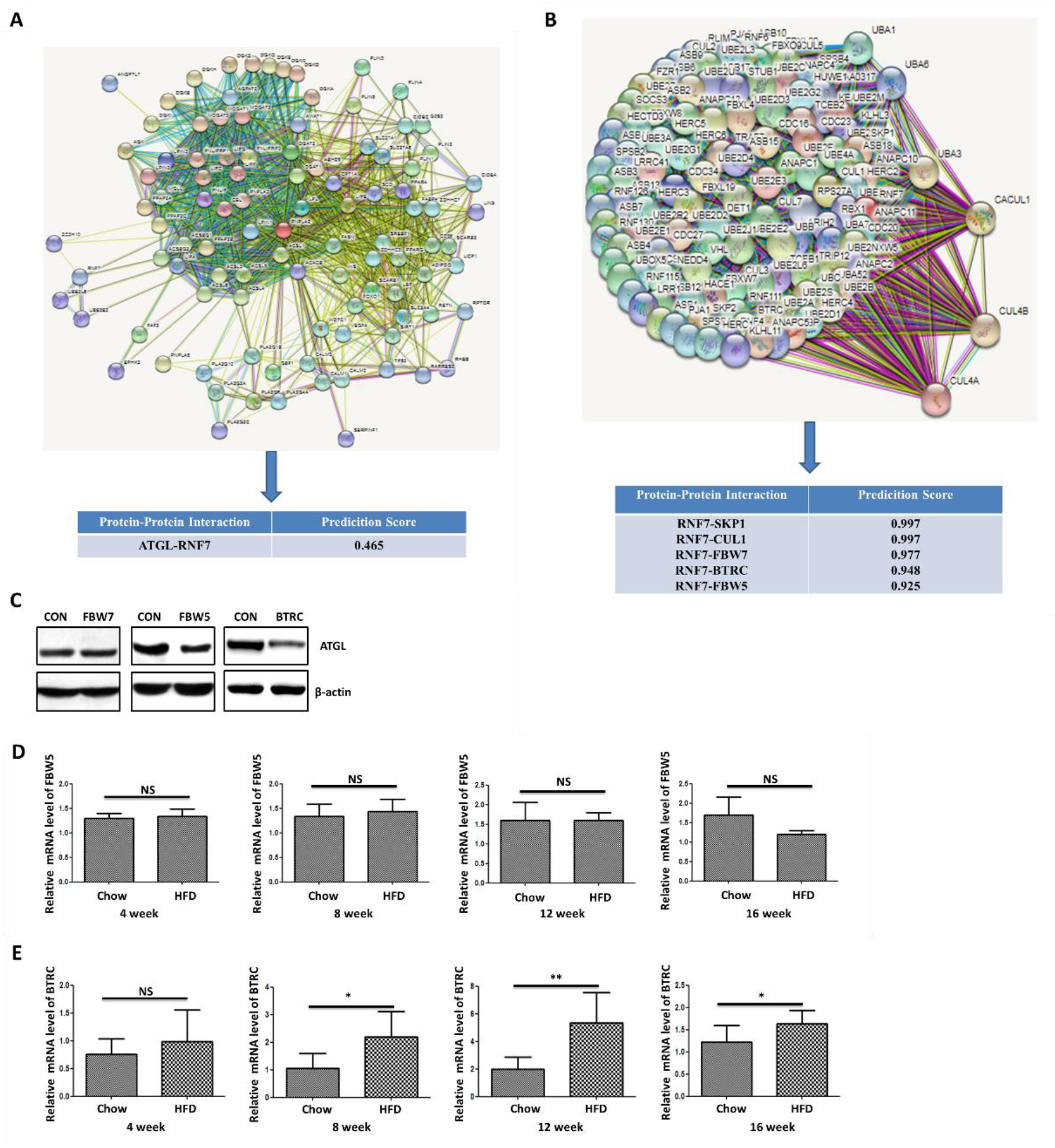
BTRC down-regulates ATGL level. A, B Bioinformatics to predict E3 enzymes that maybe bind ATGL. The STRING database (https://string-db.org/) indicated that RNF7 maybe interact with ATGL (A), SKP1, CUL1, FBW7, BTRC and FBW5 (B), and their predicted interactions scores were 0.465, 0.997, 0.997, 0.997, 0.948 and 0.925 respectively. C HepG2 cells were overexpressed with FBW7, FBW5 and BTRC plasmids respectively for 24h, followed by Western blot analysis for ATGL. D and E qRT-PCR analysis of expression of FBW5 (D) and BTRC (E) in the liver of high-fat diet mouse fed for 4 weeks, 8 weeks, 12 weeks and 16 weeks (n = 6 mice/group). Data are represented as Mean ± SEM of three independent assays. \**P* < 0.05 and \*\**P* < 0.01.

**Figure 4.**
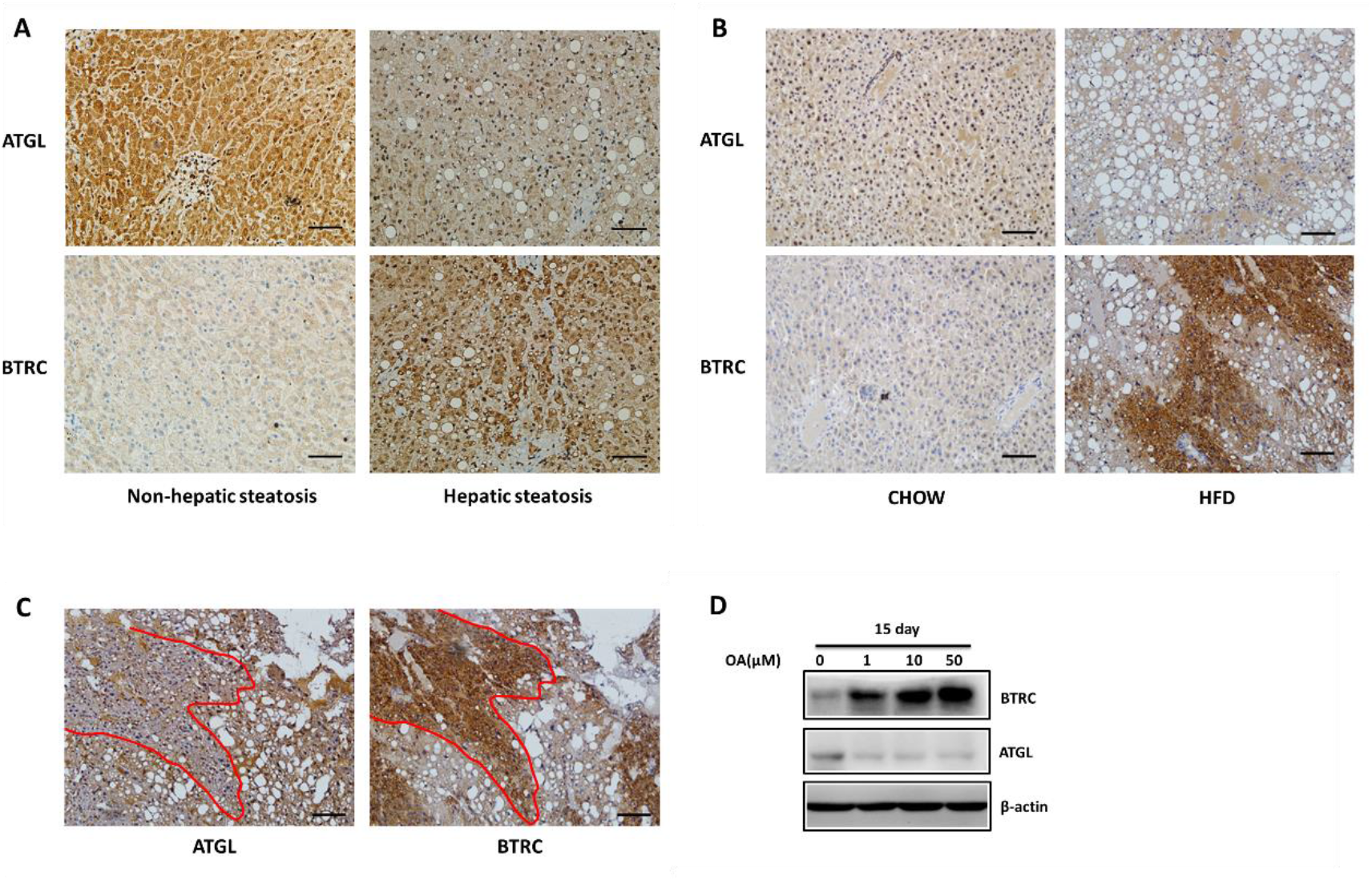
BTRC is negatively correlated with ATGL in hepatic steatosis. A, B Immunohistochemical ATGL and BTRC staining in paracancerous liver tissues of hepatic carcinoma patients with hepatic steatosis (A, n=11 samples/group), and in high-fat diet mouse fed for 16 weeks (B, n=6 mice/group). C High-fat diet mouse were fed for 16 weeks and performed IHC analysis by staining ATGL and BTRC in the same field of vision (n=6 mice/group). D HepG2 cells were treated with various concentrations of OA for 15 days. ATGL and BTRC expression was examined by immunoblotting. *Scale bars*: 100μm(A-B), 50μm(C).

### BTRC acts as the E3 Ligase of ATGL

To further identify whether BTRC acts as an E3 ligase for ATGL proteasome pathway degradation, we overexpressed BTRC in Chang liver cells. As shown in **Fig 5A**, BTRC significantly reduced ATGL protein level, while did not affect ATGL transcriptional level **(Fig 5B)**, suggesting that BTRC regulated ATGL levels at the post-transcriptional pathway rather than the transcriptional pathway. Similar results were obtained in HepG2 cells **(Fig 5C&D)**. Furthermore, BTRC overexpression resulted in a dramatic increase in ATGL ubiquitination **(Fig 5E)**. Moreover, to examine whether BTRC interacts with ATGL, we transfected ATGL and BTRC plasmids in HEK293A cells and performed a co-immunoprecipitation assay. The result showed that BTRC was bound to ATGL **(Fig 5F)**. Correspondingly, endogenous ATGL and BTRC were co-immunoprecipitated in hepatocytes and liver tissue **(Fig 5G&H)**. Taken together, our results strongly show that BTRC directly binds to ATGL and acts as an E3 ubiquitin ligase of ATGL.

**Figure 5.**
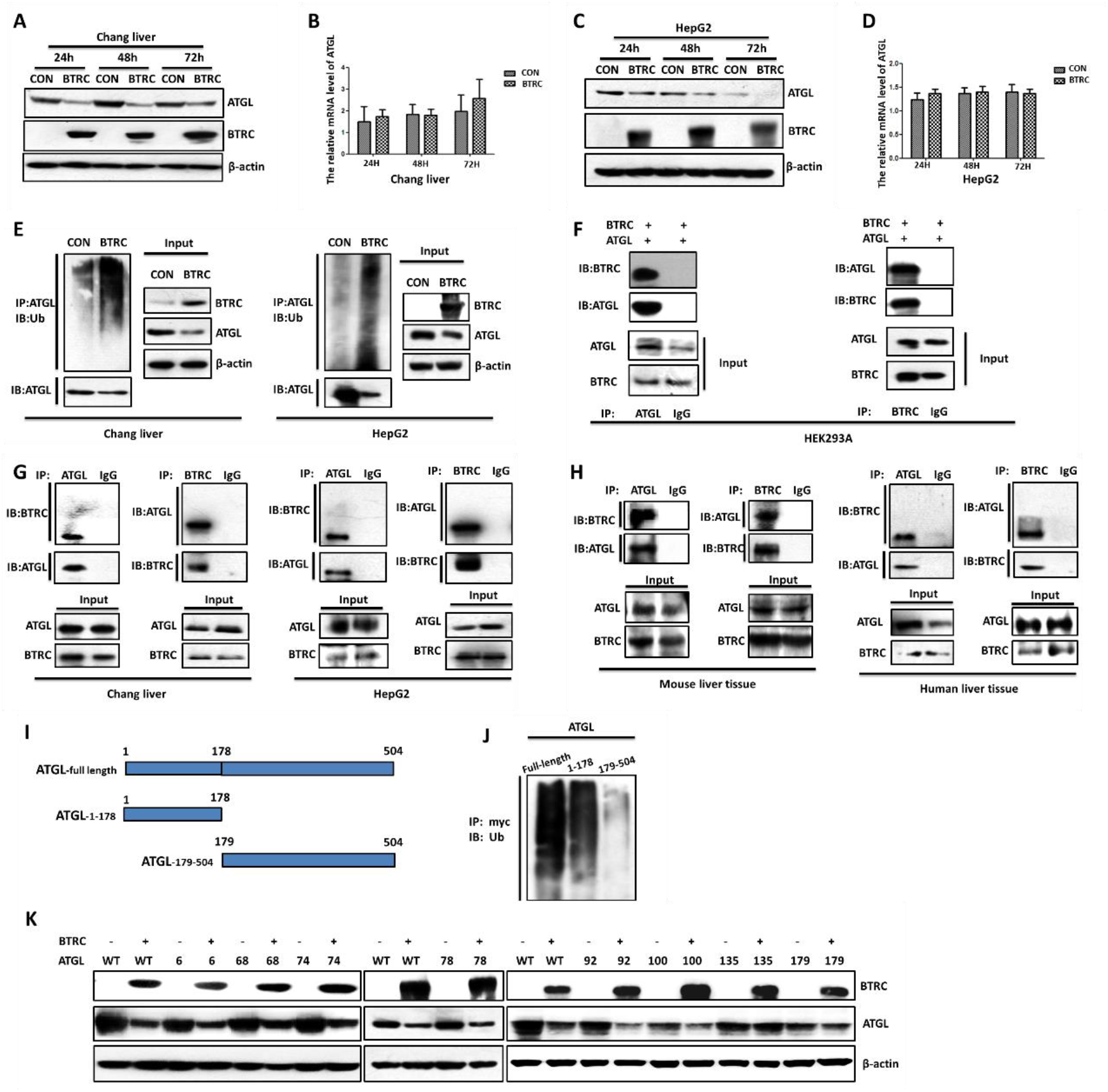
BTRC acts as the E3 ligase of ATGL. A-D Western blot and qRT-PCR analysis of ATGL protein level (A, C) and transcriptional level (B, D) in Chang liver or HepG2 cells transiently transfected with BTRC for 24h, 48h and 72h. E Cell lysates from Chang liver and HepG2 cells transiently transfected with BTRC for 48h were prepared and immunoprecipitated with anti-ATGL. The cell lysates and immunoprecipitates were analyzed by western blot with anti-Ub and ATGL. F-H The lysates form HEK293A cells co-expressed BTRC and ATGL for 24h (F), Chang liver, HepG2 cells (G), and liver tissues of mouse and patients with cholangioma (H) were performed Co-immunoprecipitation assay. Co-immunoprecipitation with anti-ATGL or anti-BTRC and ATGL and BTRC were visualized by western blotting. The co-IP assays were repeated independently three times and showed consistent results. IgG as negative control. I A graphical representation of full-length, patatin-like fragment (1–178 amino acids) and 179-504 fragment (179-504 amino acids) of ATGL. J Cell lysates from HepG2 cells transiently transfected with full-length, patatin-like fragment and 179-504 fragment of ATGL for 48h were prepared and immunoprecipitated with anti-ATGL. The cell lysates and immunoprecipitates were analyzed by western blot with anti-Ub and ATGL. K HepG2 cells transiently transfected with the plasmid of BTRC and site-specific mutation of ATGL at Lys6, Lys68, Lys74, Lys78, Lys92, Lys100, Lys135, Lys179 to Arg. ATGL protein level was evaluated by western blot.

### BTRC promotes ATGL ubiquitination degradation predominantly at lysine 135 residue

We subsequently sought to identify the lysine residues of ATGL that are ubiquitinated, which was promoted by BTRC. To achieve this goal, we examined the polyubiquitination of various ATGL domains. Full-length of ATGL, patatin-like fragment (1–178 amino acids), and 179-504 fragment (179-504 amino acids) containing myc-tagged vectors were performed to IP assay **(Fig 5I)**. The result indicated that ATGL ubiquitination occurred in the patatin-like fragment **(Fig 5J)**. We further mutated the lysine residue of the patatin-like fragment to arginine residue. As shown in **Fig 5K**, Lys→Arg mutation only at the 135^th^ position significantly eliminated the degradation of ATGL by BTRC.

### BTRC decreases lipolysis and increases lipid accumulation in the oleic acid-induced hepatic steatosis cells model by targeting ATGL proteasomal degradation

Because BTRC acts as an E3 ligase for ATGL, overexpression of BTRC should decrease the lipolysis rate, increase cellular lipid accumulation, and reduce lipid mobilization. To confirm this hypothesis, hepatocytes were transfected with BTRC plasmid and stained with neutral lipids indicator Oil Red O. In the oleic acid-induced hepatic steatosis cells model, overexpression of BTRC enhanced lipid droplets accumulation **(Fig 6A&B)**. Following this, we should verify whether ATGL is necessary for BTRC-dependent lipid metabolism, and then we transiently co-expressed BTRC and ATGL in the hepatic steatosis cells model. The result showed that co-expression of ATGL significantly reversed the effect of BTRC on lipid droplet accumulation in primary hepatocytes and HepG2 cells **(Fig 6A-D)**. Consistently, BTRC enhanced TAG content and inhibited glycerol release, while co-expression of ATGL reversed the impact of BTRC on fat metabolism **(Fig 6E&F)**. On the contrary, interfering with BTRC by siRNA, lipid droplet accumulation was reduced in the oleic acid-induced hepatic steatosis cells model **(Fig 6G&H, Fig S2A&B)**. Taken together, those results proposed that BTRC decreases lipolysis and increases lipid accumulation *via* targeting ATGL proteasomal degradation.

**Figure 6.**
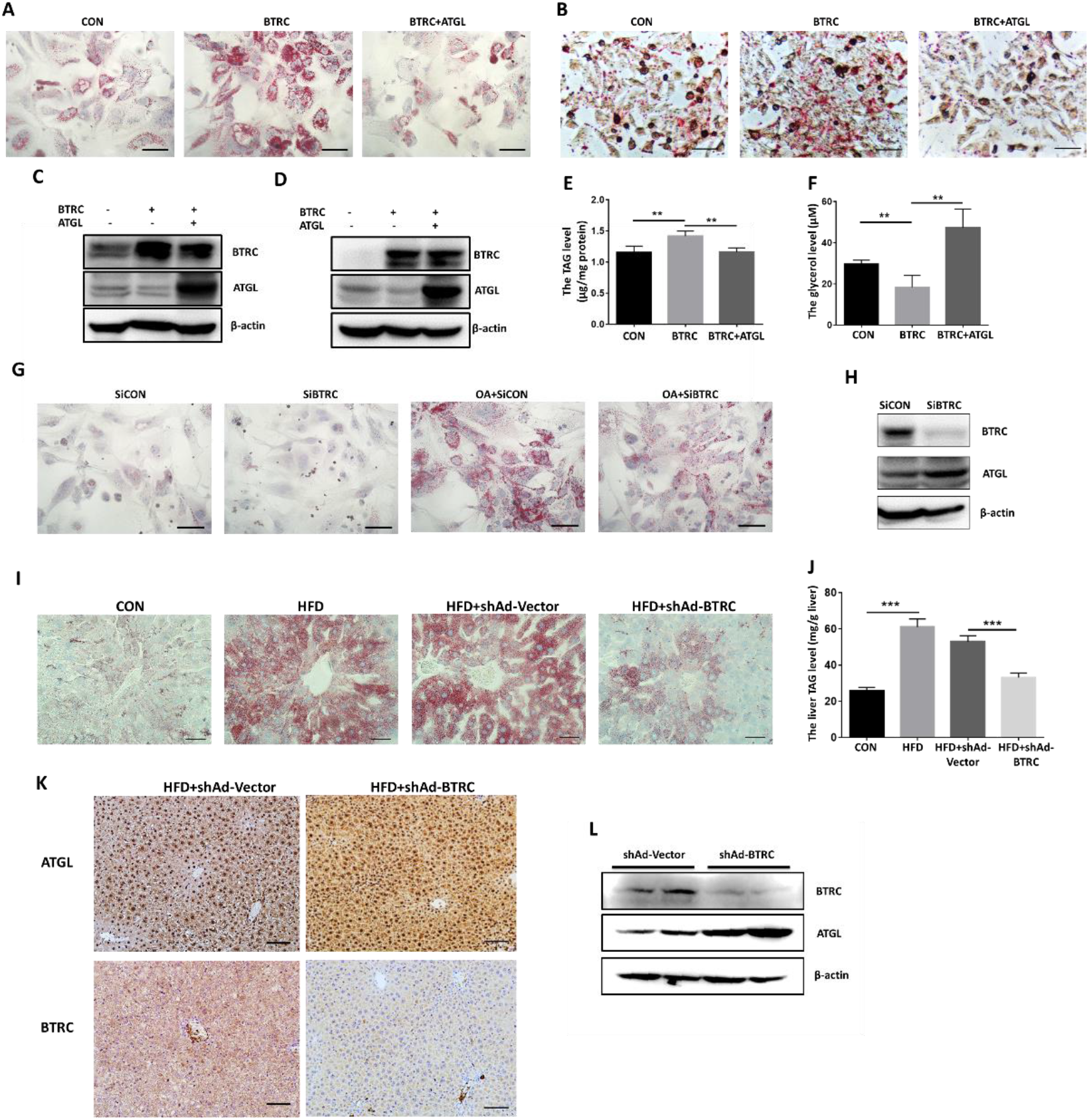
BTRC controls lipid accumulation and fat metabolism in hepatic steatosis cells model and HFD mice by targeting ATGL proteasomal degradation. A-B Primary hepatocytes (A) and HepG2 cells (B) were treated with 100μM OA for 48h to induce hepatic steatosis followed by transiently transfecting with BTRC or BTRC and ATGL plasmids. Lipid droplets were assessed by Oil Red O staining assay. C Western blot analysis of ATGL and BTRC protein level in primary hepatocytes treated with the same condition as (A). D-F Western blot analysis of ATGL and BTRC protein level (D)detection of the cellular TAG (E) and glycerol (F) concentrations by commercialized Kit in HepG2 cells treated with the same condition as (B). G, H Interfered with BTRC by siRNA for 48h in OA-induced hepatic steatosis primary hepatocytes followed by Oil Red O staining (G) and ATGL and BTRC detecting by Western blot analysis (H). I, J After six weeks of high-fat-diet feeding, the tail vein was injected with adenovirus specifically knockdown BTRC (shAd-BTRC) for three weeks (n = 5 mice/group). Lipid droplets accumulation analysis by Oil Red O staining (I, *Scale bars*: 50μm), TAG level in liver(J) detected by commercialized Kit. K, L Expression of ATGL and BTRC were detected with immunohistologic staining (K) and western blot (L) in injecting shAd-BTRC animals. Data are represented as Mean ± SEM of three independent assays. \**P* < 0.05, \*\**P* < 0.01; \*\*\**P* < 0.001. *Scale bars*: 50μm (A, B, G, I) and 100μm (K).

### BTRC knockdown ameliorates hepatic steatosis in HFD-mice

Next, we examined the effect of BTRC in the HFD-induced hepatic steatosis mouse model. After six weeks of HFD-fed, the tail vein was injected with adenoviruses to knockdown BTRC (shAd-BTRC) for three weeks. Compared with adenovirus vector (shAd-Vector)-injected mice, lipid droplet accumulation and TAG levels of the liver were expectedly lower in shAd-BTRC-injected mice **(Fig 6I&J)**, besides a significant reduction of BTRC expression with a noticeable increase of ATGL protein levels in the liver **(Fig 6K&L)**. Nonetheless, no corresponding alteration of FFA and TAG was observed in the plasma of shAd-BTRC-injected mice **(Fig S3A&B)**. Similar results were also tested in another way of animal treatment (Injection of adenovirus and HFD simultaneously for four weeks, **Fig S4A-C**). Collectively, our result demonstrates that knockdown of BTRC increases ATGL levels, inhibits TAG accumulation, and ameliorates hepatic steatosis in the HFD-induced fatty liver mice.

## Discussion

The present study uncovers a pivotal pathogenic role of BTRC in developing NAFLD. Such a point of view is supported by the fact that BTRC *in vivo* and *in vitro* could attenuate lipolysis and contribute to the development of hepatic steatosis, potentially *via* downregulating the expression of ATGL. In cholangioma patients, hepatic carcinoma patients, and obese mice models, we demonstrated that proteasomal degradation of hepatic ATGL resulted in the ATGL decrease, which was positively associated with increased hepatic lipid droplet accumulation. In the human liver cell model, we identified BTRC as an E3 ligase for ATGL, which was mediated by polyubiquitination of patatin-like fragment predominantly at lysine 135. Moreover, knockdown of BTRC could ameliorate hepatic steatosis *via* inhibiting ATGL proteasomal degradation and upregulating ATGL expression, which might be a new therapeutic target of NAFLD.

NAFLD patients with insulin resistance had higher liver steatosis grades and lower ATGL expression^12^. Our results consistently showed that ATGL protein was significantly down-regulated in cholangioma patients with hepatic steatosis, hepatic carcinoma patients with hepatic steatosis, hepatic steatosis of mice induced by HFD, and oleic acid-induced hepatic steatosis cells model **(Fig 1)**. Protein expression is generally regulated at the transcriptional or post-transcriptional level. Our results showed the transcription level of ATGL remained unchanged and the protein level of ATGL decreased in hepatic steatosis of mice induced by HFD and oleic acid-induced hepatic steatosis cells model **(Fig1D-G)**, which makes us focus on the post-transcriptional regulation. The proteasome and lysosome pathways are the primary forms of post-transcriptional level regulation in cellular protein turnover^15^. Our previous study reported that ATGL could be degraded in adipocytes in a ubiquitin-dependent proteasome way, which was promoted by Pigment Epithelium-Derived Factor (PEDF)^25^, while it is unclear in hepatocytes. Our data showed that the amount of ATGL gradually decreased as time went on under CHX treatment (Fig 2B&D), as well MG132 could block its degradation in hepatocytes **(Fig 2A, 2C&E)** and hepatocytes with fatty degeneration **(Fig 2F)**. These results indicate that ATGL degradation is regulated by the proteasome-mediated degradation pathway in hepatocytes and hepatic steatosis. To further clarify the mechanisms involved in ATGL degradation, we focus on the E3 ligase targeting for ATGL.

The Skp1–cullin–F-box protein (SCF) family of ubiquitin E3 ligases complexes is composed of S-phase kinase-associated protein 1 (Skp1), cullin 1 (Cul1), a variable member of F-box family proteins, and RBX/ROC RING component (RBX1/ROC1 or RBX2/ROC2/SAG/RNF7)^20,24^. SCF complexes are the most prominent family of E3 ligases that ubiquitinate various regulatory proteins for 26S proteasome degradation, which have been shown to regulate multiple biological processes, such as apoptosis, development, lipid metabolism, etc^20,23^. BTRC, one of the F-box proteins, also called β-TrCP, has been reported to be the E3 ligase of many essential molecules, including IκBα, and β-catenin, which was considered to be related to cancer development and inflammation^20,26^. It is reported that another F-box protein, FBW7, could promote the degradation of the SREBP family of transcription factors and regulate lipogenesis^27,28^. Our results identify that BTRC rather than FBW7 is the E3 ligase for ATGL. Recently, Mainak Ghosh, et al. reported that COP1(also known as RFWD2), which acted as an E3 ligase, could promote hepatic TAG accumulation by promoting ATGL degradation^29^. PEDF diminishes ATGL protein stability by promoting its proteasomal degradation in a COP1-dependent manner^30,31^. Nevertheless, COP1 promotes ATGL ubiquitination predominantly on the 100^th^ residue rather than the 135^th^ residue regulated by BTRC **(Fig 5K)**, suggesting that BTRC is a new E3 ligase for ATGL besides COP1. Moreover, although adenovirus-mediated depletion of COP1 ameliorates HFD-induced steatosis in mouse liver and improves liver function, Mainak Ghosh, et al. did not measure whether COP1 was up-regulated in hepatic steatosis in patient samples and animal models with hepatic steatosis; the above researches revealed that the role of COP1 in the development of hepatic steatosis is uncertain. Thereby, unraveling the function of BTRC as a new E3 ligase for ATGL and its pathophysiology role in the development of NAFLD is pivotal for us.

In various hepatic steatosis animal models and human hepatic steatosis tissues, the protein level and mRNA level of BTRC are increased, positively associated with lipid droplet accumulation **(Fig 3E&4)**. Furthermore, overexpression of BTRC could promote the degradation of ATGL **(Fig 5)**. Conversely, knockdown of BTRC *in vitro* and *in vivo* could increase ATGL expression, resulting in inhibiting TAG accumulation and ameliorating hepatic steatosis **(Fig 6)**. In short, our study demonstrated that up-regulated BTRC in the liver promoted ATGL degradation, followed by hepatic steatosis. The spectrum of NAFLD ranges from simple steatosis to NASH, followed by cirrhosis, and a subset of the latter will finally advance to hepatocellular carcinoma; inflammation runs through and promotes the disease processes^32^. Besides, BTRC is likewise an E3 ligase for IκBα, degradation of which would induce NF-κB translocation to the nucleus, then enhance the expression of target genes encoding many inflammatory mediators causing the development of inflammation^33^. In addition to ATGL degradation, the critical role of continuously elevated BTRC on liver inflammation is worth further investigation.

In conclusion, we identify BTRC, as a novel E3 ligase for ATGL, could decrease ATGL expression *via* mediating its ubiquitin-dependent proteasome degradation at lysine 135 of patatin-like fragment, resulting in the hepatic TAG accumulation and development of hepatic steatosis. Knockdown of BTRC increases ATGL levels, inhibits TAG accumulation, and ameliorates hepatic steatosis. Our finding has translational therapeutic implications for tackling and reversing NAFLD in humans.

## Author contributions

Guoquan Gao and Weiwei Qi designed and performed experiments, analyzed data, and wrote the manuscript. Xia Yang and Yong Xia designed and performed experiments and wrote the manuscript. Zhenzhen Fang, Chuanghua Luo, Yongcheng Zeng and Yanlan Long performed experiments and assessed the outcome. Honghai Hong provided human samples and designed experiments. Zhiyu Dai designed experiments. Ti Zhou and Junxi Liu designed experiments, analyzed data, and wrote the manuscript. All authors reviewed and commented on the manuscript.

## Supplemental materials

### Plasmids and Vectors

Human BTRC, FBW7, FBW5 and ATGL clones were constructed using Ruyilian Kit from SiDanSai Biotechnology (Sidansai, Shanghai, China) according to the manufacturer’s instruction. In briefly, human BTRC, FBW7, FBW5 and ATGL were cloned from cDNA using specific primiers (Table S1), and then fused to the vector provided by Ruyilian Kit. SiRNA for BTRC was purchased from Ribobio Company (Guangzhou, China). The sequence of the BTRC siRNA was 5’-GACTACAGTTTGATGAATT-3’(for human); 5’-GCGACATAGTTTACAGAGA-3’(for mouse). Cells were transfected with the BTRC siRNA using Lipofectamine 2000 (Invitrogen, Carlsbad, CA, USA) according to the manufacturer’s instructions. Mutagenesis of the ATGL clone was done using In-Fusion technology from Clontech Company (Catalog 639650, Clontech, Japan) according to the manufacturer’s instruction. In briefly, ATGL lysine site mutation fragment was cloned from ATGL plasmids using specific mutant primiers (Table S2), and then connected with the linear vector cloning using corresponding primers (Table S2).

**Table S1.**
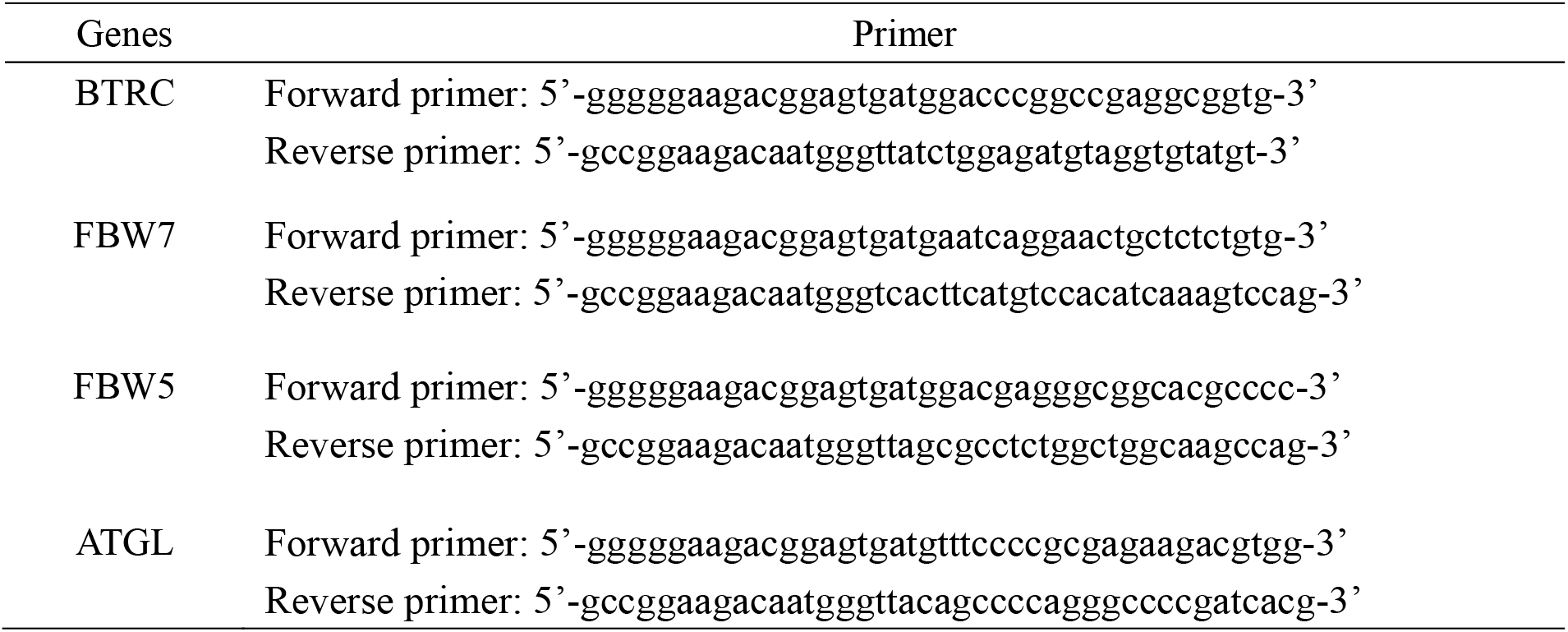
The primers for cloning Human BTRC, FBW7, FBW5 and ATGL using Ruyilian Kit Genes Primer

**Table S2.**
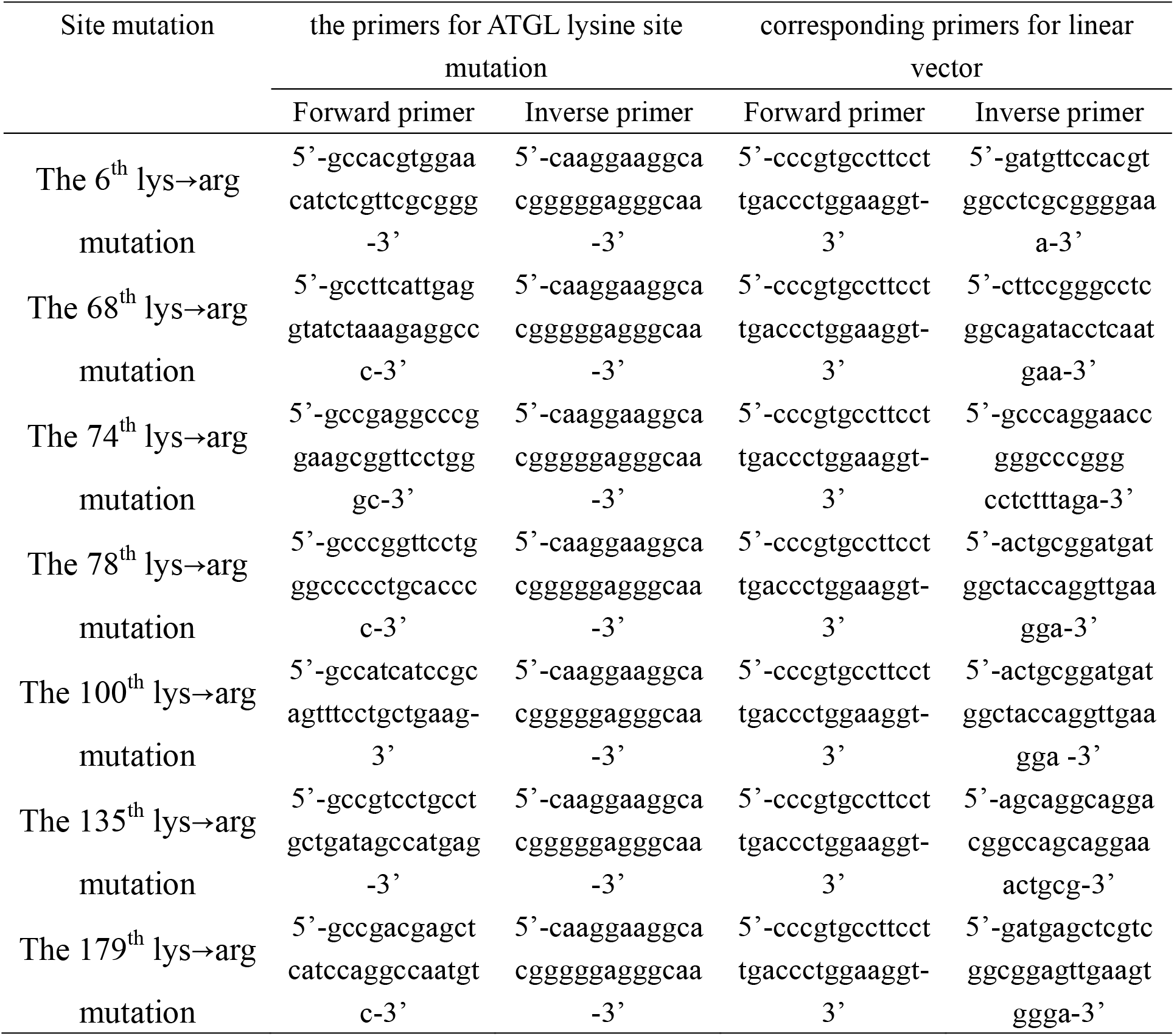
The primers for ATGL lysine site mutation and corresponding primers for linear vector

## Supplemental Figure

**Figure S1.**
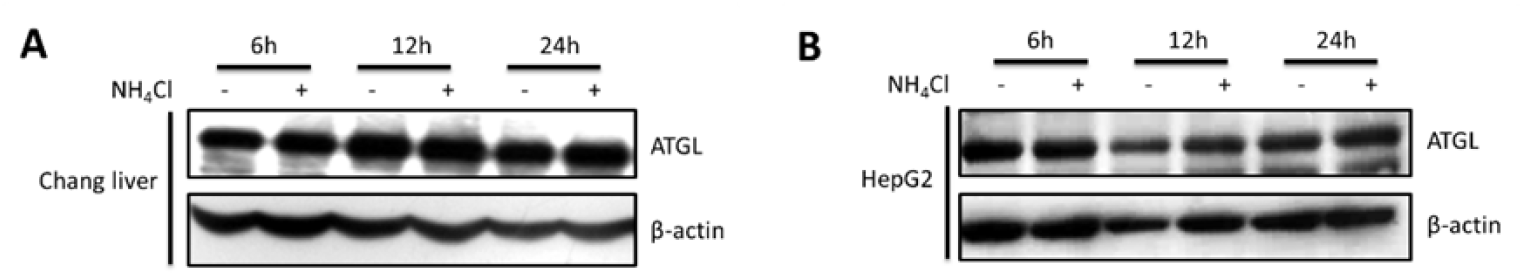
The level of ATGL in Chang liver and HepG2 cells with the treatment of lysosomal inhibitor ammonium chloride. A, B ATGL was detected by immunoblotting in Chang liver cells (A) and HepG2 cells (B) treated with ammonium chloride for indicated time. β-actin was used as a loading control. NH_4_Cl: 10mM. Data are represented of at least 3 independent experiments.

**Figure S2.**
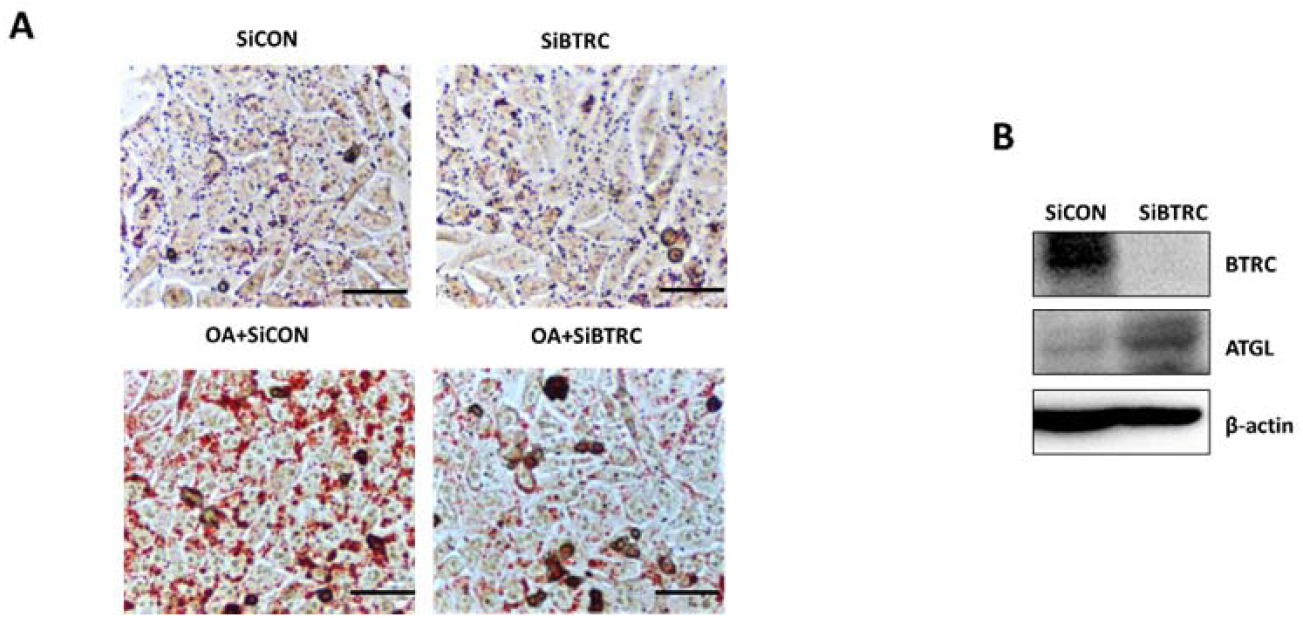
Interfering with BTRC by siRNA, lipid droplet accumulation was reduced in the oleic acid-induced HepG2 cells. A, B Interfered with BTRC by siRNA for 48h in OA-induced hepatic steatosis HepG2 cells followed by Oil Red O staining (A) and ATGL and BTRC detecting by Western blot analysis (B). *Scale bars*: 50μm.

**Figure S3.**
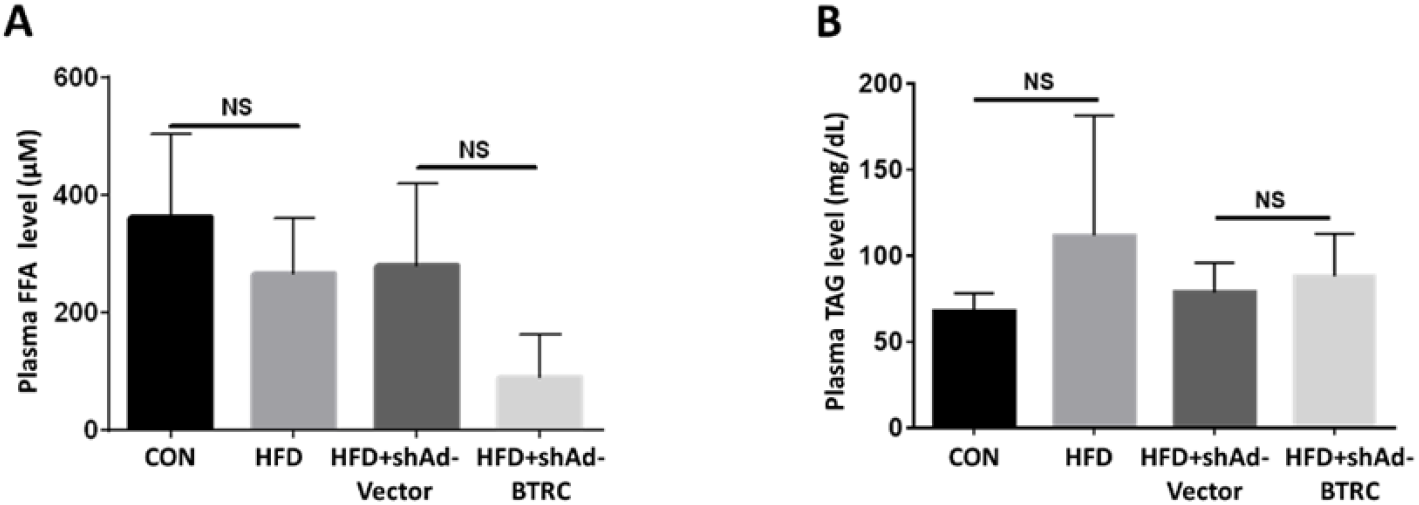
FFA and TAG level in plasma of injecting shAd-BTRC animals. A, B After six weeks of high fat feeding mice, the tail vein was injected with adenovirus specifically knockdown BTRC (shAd-BTRC) for three weeks. FFA level (A) and TAG level (B) in plasma detected by commercialized Kit in injecting shAd-BTRC animals. Data are represented as Mean ± SEM of three independent assays.

**Figure S4.**
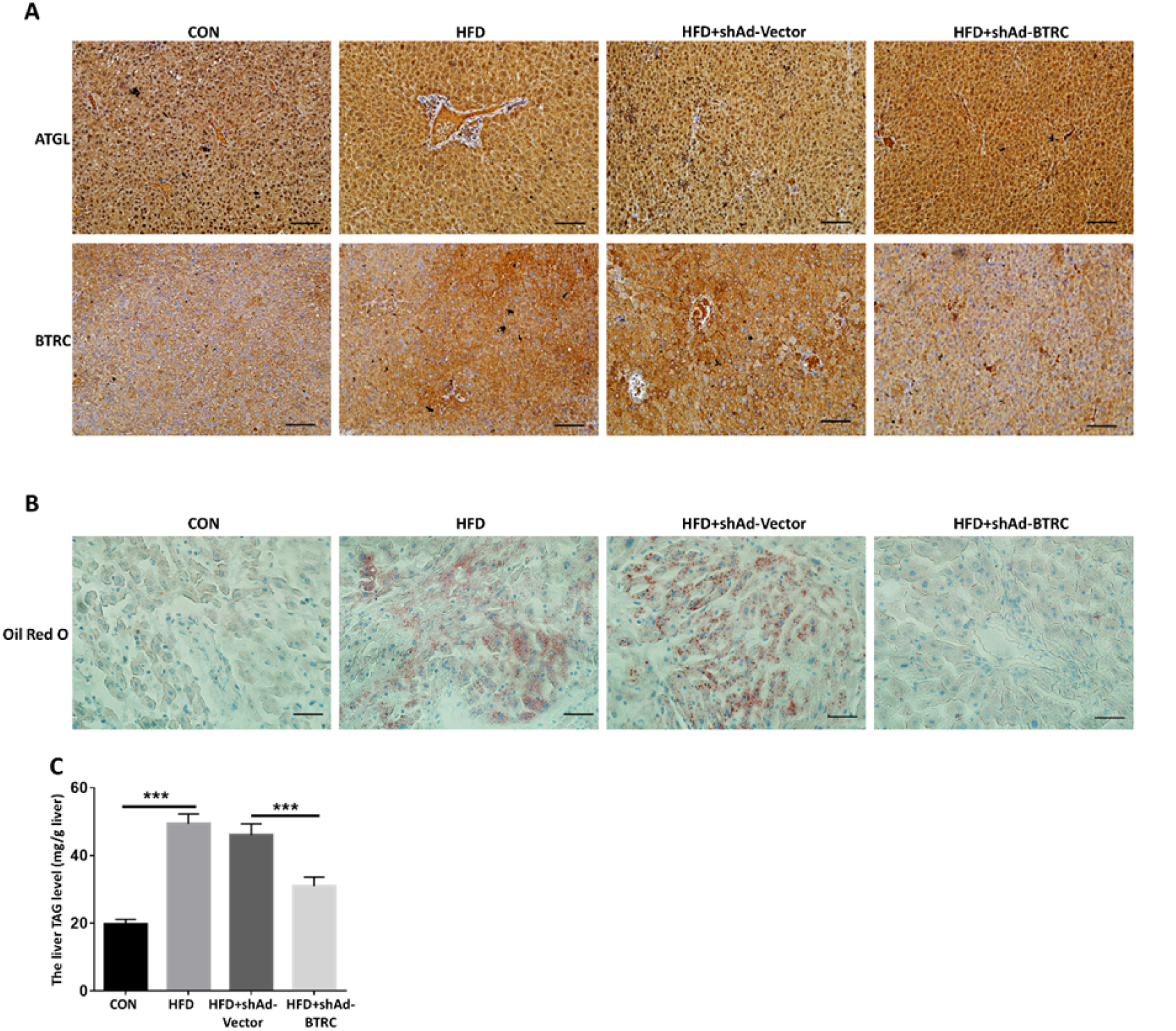
BTRC knockdown ameliorates hepatic steatosis in high-fat diet mouse. A, B The mice were injected shAd-BTRC adenovirus and high-fat diet for four weeks simultaneously, followed by assessment of ATGL and BTRC expression by immunohistologic staining, and lipid droplets accumulation analysis by Oil Red O staining. C TAG lever in live was detected by commercialized Kit in injecting shAd-BTRC animals. Data are represented as Mean ± SEM of three independent assays. \*\**P* < 0.01; \*\*\**P* < 0.001. *Scale bars*: 100μm (A) and 50μm (B). n = 5 mice/group.

